# Dexamethasone impairs glycolysis but improves mycobacterial killing in primary human macrophages

**DOI:** 10.1101/2025.06.27.661762

**Authors:** Lorraine Thong, Olivia Sandby Thomas, Oisin Ó Gallchobhair, Emily Duffin, Anjali S Yennemadi, Gina Leisching, Dearbhla M Murphy, Parthiban Nadarajan, Finbarr O’Connell, Mary P O’Sullivan, Sharee A Basdeo, Donal J Cox, Joseph Keane

## Abstract

Glucocorticoids (GC) are useful adjunctive host directed therapies for sub-types of tuberculosis (TB). Macrophages play a central role in controlling *Mycobacterium tuberculosis* (Mtb) infection, relying on glycolytic reprogramming to support an effective host defense, yet the influence of GC on these important phagocytes is poorly understood. Here, we examined the impact of dexamethasone on metabolic and functional responses of primary human airway macrophages (AM) from bronchoalveolar lavage fluid and monocyte-derived macrophages (MDM). We found that dexamethasone significantly reduced basal and compensatory glycolysis in both AM and MDM, and decreased expression of the glycolytic enzyme PFKFB3. Oxidative metabolism was lower in dexamethasone AM but not MDM, indicating different specific metabolic sensitivity of macrophages. Dexamethasone also inhibited the glycolytic response to Mtb and reduced secretion of IL-1β, TNF, IL-6, IL-8, and IL-10. Dexamethasone-treated macrophages showed enhanced survival following Mtb infection and these cells had a significant reduction in bacterial burden. This antimicrobial effect was impaired when macrophages were pre-treated with bafilomycin A1, implicating that phagosomal acidification may at least in part mediate dexamethasone-induced bacterial control. Collectively, these findings demonstrate that dexamethasone reprograms human macrophage metabolism toward a less glycolytic state while preserving their ability to limit Mtb growth. These results may offer a basis for the clinical benefit of GC in some TB presentations and support the development of targeting GC therapies to macrophages, thereby mitigating inflammation without compromising host antimicrobial defense.

## Introduction

Despite their immunosuppressive features, glucocorticoids (GC) are increasingly used as adjunctive host directed therapy for pneumonia. GC, such as dexamethasone, potently inhibit inflammatory processes in both innate and adaptive immune cells and also interfere with the metabolic functions of immune cells (1–3). The COVID-19 pandemic renewed interest in the clinical utility of dexamethasone as it was the first therapeutic intervention that improved patient survival with severe respiratory failure secondary to COVID-19 (4). Similarly, patients with severe community acquired pneumonia treated with GC also demonstrate improved rates of survival (5). GC, such as dexamethasone, are the only host-directed adjunctive treatment recommended in combination with antimicrobial regimens by the World Health Organization for tuberculosis (TB). Yet thus far only patients with TB meningitis or TB pericarditis have shown a survival benefit from adjunctive GC use (6–8). Indeed, prednisolone therapy was associated with improved clearance times of *Mycobacterium tuberculosis* (Mtb) from sputum of HIV positive patients (9).

Airway macrophages (AM) play a critical role at the frontline of immunity, functioning as key regulators of inflammation and defending against pathogens, such as Mtb. The metabolic state of macrophages profoundly influences their activity, with inflammatory macrophages exhibiting increased glycolysis which supports functions such as increased cytokine production and antimicrobial responses (10). Recent studies examining the effect of dexamethasone on cellular metabolism have thus far focused on examining the response to lipopolysaccharide (LPS) utilizing modified mitochondrial and glycolytic stress tests (1–3). There is a lack of data on the metabolism and downstream effector functions of primary human macrophages when treated with GC in response to mycobacterial infection.

We therefore investigated the effect of dexamethasone on glycolytic metabolism in human tissue resident airway, and peripheral blood derived macrophages. These data demonstrate that dexamethasone reduces glycolytic metabolism in both tissue resident AM and monocyte derived macrophages (MDM), and impairs the glycolytic shift induced by mycobacterial stimulation. Furthermore, dexamethasone treated Mtb infected macrophages had improved survival which was associated with reduced secretion of both pro- and anti-inflammatory cytokines. Dexamethasone also improved the ability of the human macrophage to limit mycobacterial growth. These data indicate that while dexamethasone can limit inflammation it does not adversely impact mycobacterial control in human macrophages.

## Methods

### Human Macrophage Culture

Buffy coats were obtained with consent from healthy donors (aged between 18–69; ethical approval, School of Medicine Research Ethics Committee, Trinity College Dublin). Peripheral blood mononuclear cells (PBMC) were isolated by density-gradient centrifugation over Lymphoprep (StemCell Technologies). Cells were resuspended in RPMI (Gibco) supplemented with 10% AB human serum (Merck) and plated onto non-treated tissue culture plates (Costar) for 7 days. Non-adherent cells were removed by washing every 2-3 days. Cultures were >90% pure based on co-expression of CD14 and CD68.

Human AM were retrieved at bronchoscopy, (ethical approval, the SJH/TUH Joint Research Ethics Committee) previously as reported (11). AM were plated in RPMI (Gibco) supplemented with 10% FBS (Gibco), fungizone (2.5 μg/ml; Gibco) and cefotaxime (50 μg/ml; Melford Biolaboratories). Cells were incubated for 24 h before washing to remove non-adherent cells. Adherent cells (predominantly AM) were used for experiments.

### Macrophage Stimulation

Human macrophages were left untreated or treated with vehicle (ethanol, 0.02%) or dexamethasone (0.05-5 μM, as indicated). To model pathways of cellular death, MDM were treated with staurosporine (1 μM; Tocris) for 24.

### Macrophage Infection

Mtb H37Ra was obtained from The American Type Culture Collection (ATCC 25177™; Manassas, VA) and propagated in Middlebrook 7H9 medium supplemented with ADC (Beckton Dickinson), to log phase. Irradiated H37Rv (iH37Rv) was gifted by BEI Resources. The multiplicity of infection (MOI) and donor variation in phagocytosis of Mtb was adjusted for by Auramine-O staining, as previously described (10, 12). MDM or AM were plated on 8-well Lab-Tek chamber slides (Nunc) and infected with a range of bacterial concentrations for 3 h before extracellular bacteria were thoroughly washed off. Cells were fixed with 2% PFA, stained with Hoechst 33342 (10 μg/ml; Merck), and rapid Auramine O staining set (Scientific Device Laboratory Inc). The numbers of bacilli per cell were counted in at least 30 fields of vision per well on a fluorescent microscope (Olympus IX51).

### Metabolic Flux Analysis

MDM were placed in ice-cold PBS and incubated at 4 °C on ice for 30 min, then gently scraped and counted using trypan blue. MDM (1x10^5^ cells/well) were re-plated onto 24 well Seahorse plates (Agilent), as previously described (13). AM (1x10^5^ cells/well) were directly plated onto Seahorse plates and washed after 24 h. Glycolytic rate assays and Mitostress tests were conducted as per manufacturer’s protocol.

### Cytokine Assays

IL-1β, IL-6, IL-8, IL-10 (BioLegend), and TNF (Invitrogen) concentrations in supernatants were quantified by ELISA, according to manufacturer’s protocol.

### Gene Expression Analysis

RNA extractions from monocytes were performed using an RNeasy Mini Kit (Qiagen) according to the manufacturer’s instructions. RNA content and quality were quantified and assayed, respectively, using a Nanodrop (Thermo Fisher Scientific) and RNA reverse transcribed using the SensiFast Reverse-Transcription Kit (Meridian Biosciences). Catalogued TaqMan (Thermo Fisher Scientific) predesigned gene primer probes attached to the FAM dye were used: PFKFB3 (Hs00998698_m1), GAPDH (Hs02786624_g1), PKM2 (Hs00761782_s1), and ATP5B (Hs00969569_m1). 18S (Hs03003631_g1) was used as the reference gene primer, attached to the VIC dye. RT-qPCR was performed using SensiFast Probe Mix (Meridian Biosciences, catalog BIO-82005) on a QuantStudio 5 RT-qPCR System (Applied Biosystems). Relative quantitative data were obtained and analyzed utilizing the 2–ΔΔCt method.

### PI inclusion Assay

Human MDM were stained with propidium iodide (PI; 5 μg/ml; Merck), and Hoechst 33258 and 33342 (both 50 μg/ml; Merck) at the indicated time-points post-infection with Mtb or treatment with staurosporine or TNF. Cells were incubated for 30 min at room temperature in the dark. Imaging was performed with a LionHeart FX® Automated Microscope (Agilent) at 10X. The ratio of PI–positive cells to the total number of Hoechst 33258/Hoeschst 33242-stained nuclei in 4 fields was determined by the Gen5 software (Biotek) and used to calculate percentage cell death.

### Caspase Assay

MDM were stained with CellEvent™ Caspase 3/7 Green Detection Reagent (Invitrogen) as per manufacturer’s instructions as well as Hoechst 33258 and 33342 (both 50 μg/ml; Merck). Imaging was performed using a LionHeart FX® Automated Microscope (Agilent) at 10X. The ratio of caspase 3/7– positive cells to the total number of Hoechst 33258/Hoeschst 33342-stained nuclei in 4 fields was determined by the Gen5 software (Biotek) and used to calculate percentage Caspase 3^+^/7^+^ MDM.

### Bacterial Growth Assays

Colony forming units (CFU) were determined at 3, 48 or 120 h post-infection, as indicated. MDM were also treated with bafilomycin A1 (50 nM) or rapamycin (250 ng/ml) to examine the role of autophagy in dexamethasone induced bacterial control. Cells were lysed with triton-X 100 (0.1%) and pooled with bacterial pellets (except 3 h) from the centrifugation of supernatants. Bacteria were diluted in Middlebrook 7H9 broth and plated onto Middlebrook 7H10 agar supplemented with OADC (both Becton Dickinson) and cycloheximide (Merck). Agar plates were incubated at 37°C for 21 days and CFU were then enumerated.

### Statistical Analysis

Statistical analyses were performed using GraphPad Prism 10. Statistically significant differences between three or more groups were determined by one-way ANOVA with an appropriate multiple comparisons tests as stated. Statistically significant differences between two or more groups containing more than one variable were determined by two-way ANOVA with an appropriate multiple comparisons tests as stated. p-Values of ≤0.05 were considered statistically significant and denoted with an asterisk. Alternatively, p-values of ≤0.05 were denoted with a hashtag where a student’s t-test was applied.

## Results

### Dexamethasone reduces metabolism in primary human monocyte derived and airway macrophages

Recent evidence suggests that glucocorticoids, specifically dexamethasone, can rewire murine macrophage metabolism (1). Moreover, dexamethasone can impair increases in glycolysis in response to LPS in both murine BMDM and human MDM, although subtle differences in the glycolytic response were evident between the species (3). We recently reported that distinct metabolic differences exist within human macrophage populations when stimulated with cytokines or bacterial ligands (10). We therefore wanted to examine the effect of dexamethasone on metabolism, specifically glycolysis, in different human macrophage populations. In order to assess if dexamethasone altered glycolytic metabolism in human MDM, a glycolytic rate assay was performed to calculate the glycolytic proton efflux rate (GlycoPER; Figure 1A) by excluding the CO_2_ dependent acidification (i.e. mitochondrial acidification). MDM were left untreated or treated with vehicle control (VC; ethanol 0.02%) or dexamethasone (0.5 or 5 μM) for 24 h prior to a glycolytic rate assay. GlycoPER was used to derive the rates of basal and compensatory glycolysis (forced glycolysis by inhibiting oxidative phosphorylation). There was no difference in any metabolic parameter between untreated and VC treated MDM (Figure S1), therefore VC were used as controls for all subsequent experiments. Dexamethasone (0.5 or 5 μM) significantly reduced both the basal (Figure 1B) and compensatory (Figure 1C) glycolysis in MDM compared with VC treated MDM. The oxygen consumption rate (OCR) was assessed during the glycolytic rate assay and as expected rotenone and antimycin A reduced OCR confirming mitochondrial inhibition (Figure 1D), validating the induction of compensatory glycolysis in MDM (Figure 1A). Basal OCR was not changed in MDM treated with either concentration of dexamethasone compared with VC (Figure 1E). To depict shifts in MDM metabolism, energy maps were generated based on basal glycolysis and OCR (Figure 1F). MDM displayed a leftward horizontal shift, indicating a less glycolytic phenotype after treatment with dexamethasone (Figure 1F).

**Figure 1:**
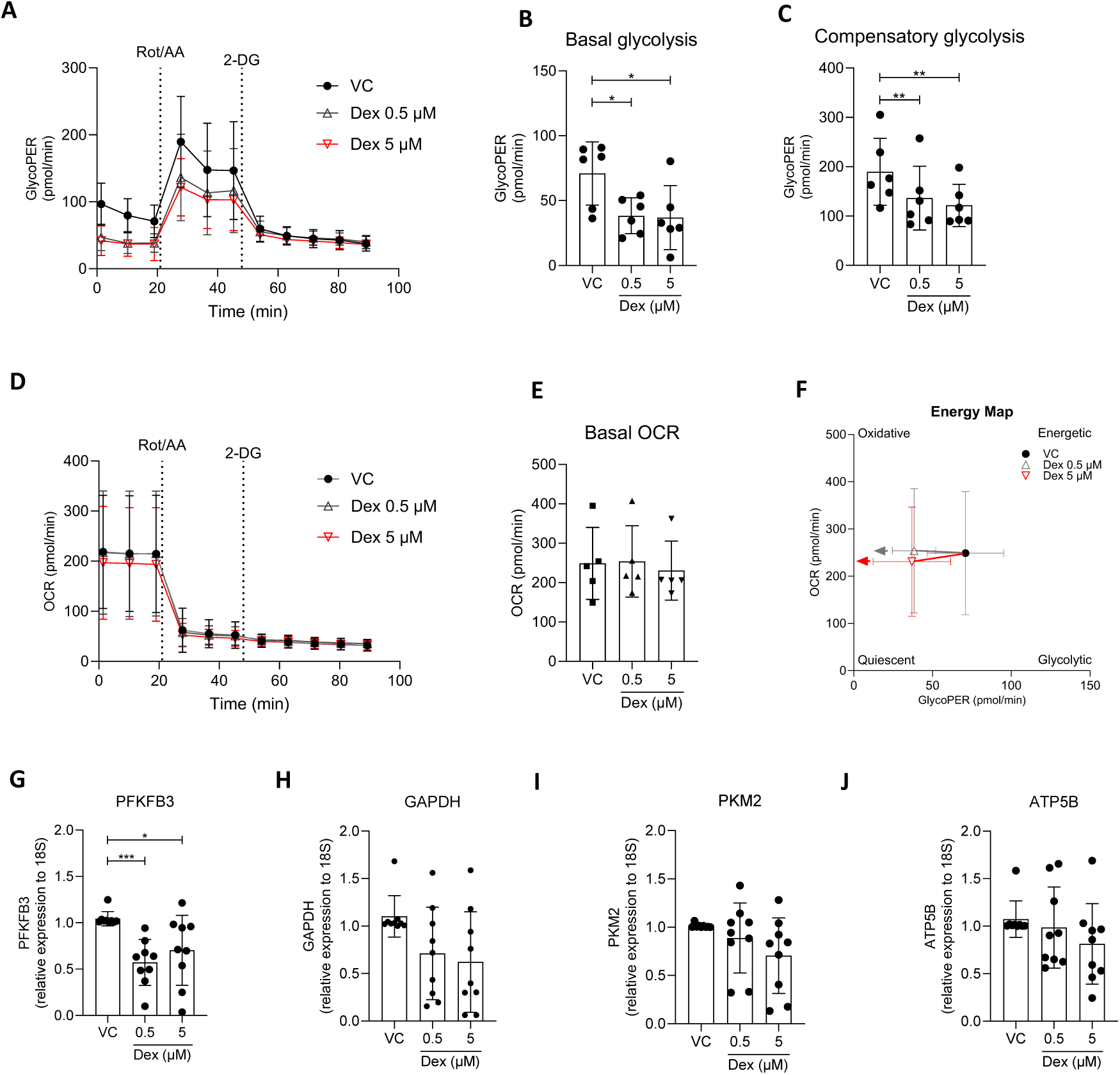
Dexamethasone reduces basal and compensatory glycolysis in human MDM. PBMC were isolated from buffy coats and MDM were adherence purified for 7 days in 10% human serum. MDM were cold lifted and seeded on Seahorse XFe24 (Agilent) culture plates (A-F). MDM were treated with either vehicle control (VC; ethanol 0.02%) or dexamethasone (Dex; 0.5 or 5 μM) for 24 h prior to analysis in the Seahorse XFe24 Analyzer (A-F). Rotenone/Antimycin A (Rot/AA) and 2-deoxyglucose (2DG) were injected at set intervals as per the glycolytic rate assay to acquire ECAR and OCR (D-E) from which GlycoPER (A) was derived to calculate basal (B) and compensatory glycolysis (C). Energy maps were generated from basal GlycoPER and the corresponding OCR value (E,F). Cell density was corrected for using Crystal Violet for Seahorse assays. Alternatively, MDM were treated with VC or dexamethasone for 4 hours, lysed and RNA extracted (G-J). Expression of PFKFB3 (G), GAPDH (H), PKM2 (I) and ATP5B (J) were assessed by real-time PCR relative to 18S. Data are represented as means ± standard deviation. Statistical significance was calculated using a one-way ANOVA with Dunnett’s post-test (A-F; n=6) or Friedman test with Dunn’s post-test (G-J; n=9); *p≤0.05, ** p≤0.005, *** p≤0.001.

To further examine if dexamethasone reduced glycolysis in MDM, the expression of rate-limiting enzymes in the glycolytic pathway and ATP5B, an enzyme in the electron transport chain of oxidative phosphorylation were examined. MDM were treated with VC or dexamethasone (0.5 or 5 μM) for 4 h and the mRNA expression of *PFKFB3*, *GAPDH*, *PKM2* and *ATP5B* was assessed by realtime qPCR. Dexamethasone significantly reduced the expression of *PFKFB3* (Figure 1G), the key rate-limiting enzyme at the beginning of the glycolytic pathway. *GAPDH, PKM2* and *ATP5B* were not significantly altered in MDM treated with dexamethasone (Figure 1H-J).

Since we have previously reported that MDM and AM have distinct metabolic responses when challenged with pathogenic stimuli (10), we next examined whether human AM responded to dexamethasone in a similar manner to MDM when treated with dexamethasone. Human AM were isolated from bronchoalveolar lavage fluid and treated with VC or dexamethasone (0.5 or 5 μM) for 24 h after which a glycolytic rate assay was performed (Figure 2A). Consistent with observations in MDM dexamethasone significantly reduced both the basal (Figure 2B) and compensatory (Figure 2C) glycolysis in AM compared with VC. OCR was monitored during the glycolytic rate assay to confirm mitochondrial inhibition (Figure 2D). In contrast to MDM, basal OCR was significantly reduced in the human AM by dexamethasone (Figure 2E). Basal glycolysis and OCR were plotted to demonstrate the metabolic shift in AM. AM treated with dexamethasone displayed a more quiescent metabolic phenotype compared with controls, as indicated by the movement of data points towards the origin on the energy map (Figure 2F).

**Figure 2:**
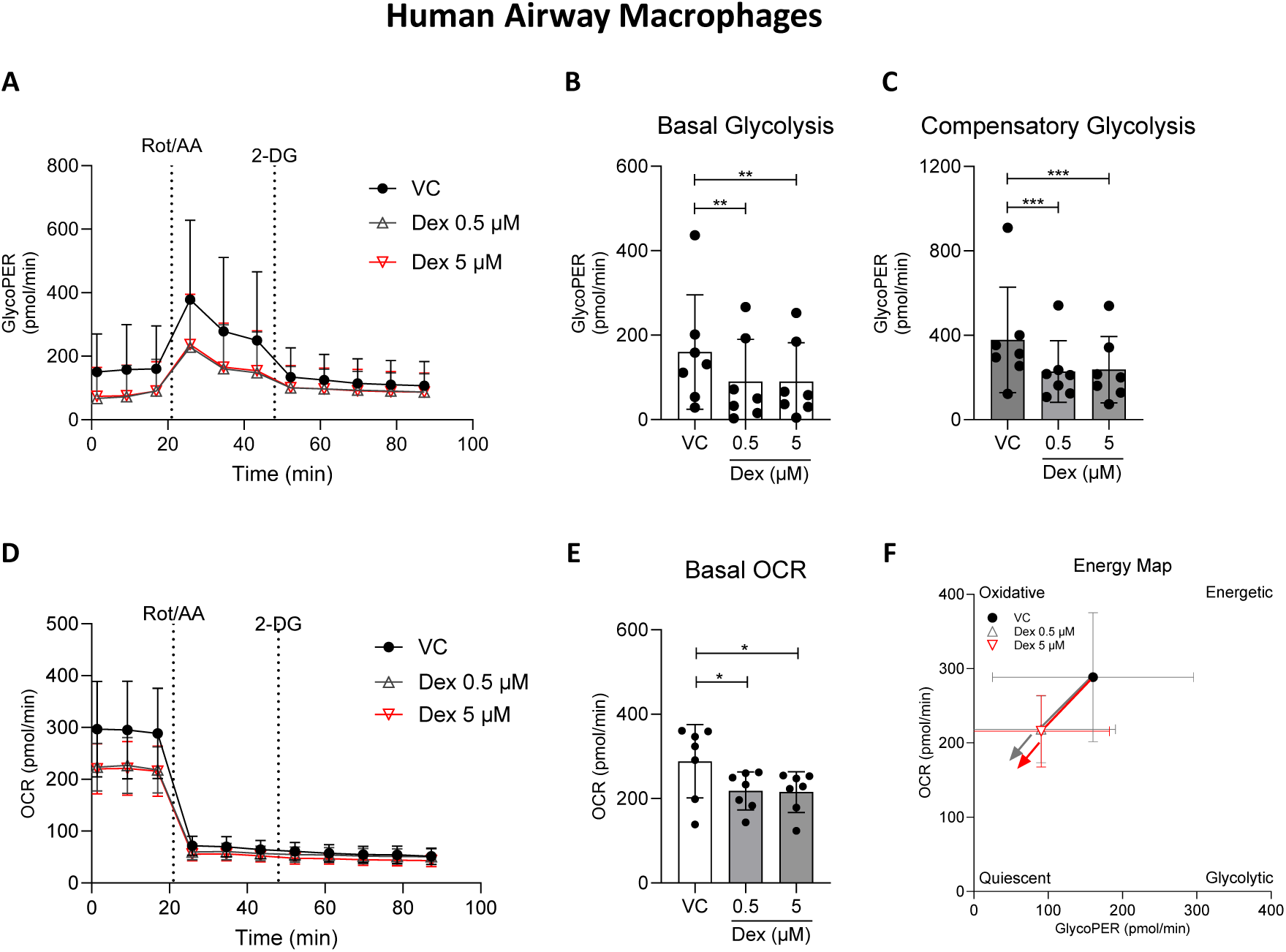
Dexamethasone reduces glycolysis and oxidative metabolism in human AM. Human AM were isolated from bronchoalveolar lavage fluid and plated directly on Seahorse culture plates. AM were treated with either VC (ethanol 0.02%) or dexamethasone (Dex; 0.5 or 5 μM) for 24 hours prior to analysis in the Seahorse XFe24 Analyzer (A-F). Rotenone/Antimycin A and 2DG were injected as per glycolytic rate assay and ECAR and OCR were acquired (D-E) from which the GlycoPER (A) was derived to calculate basal (B) and compensatory glycolysis (C). Energy maps were generated from basal GlycoPER and OCR (F). Cell density was corrected for using Crystal Violet. Data are represented as means ± standard deviation. Statistical significance was calculated using a one-way ANOVA with Dunnett’s post-test (A-F; n=7); *p≤0.05, ** p≤0.005, *** p≤0.001.

### Dexamethasone impairs the glycolytic response of primary human macrophages to Mtb

Human macrophages rapidly respond to bacteria and bacterial components such as Mtb and LPS by increasing glycolysis (13–16), to redirect cellular energy to produce inflammatory and anti-microbial mediators. Dexamethasone inhibits the glycolytic response in LPS stimulated murine and human macrophages (3). Since we demonstrated that dexamethasone reduced glycolysis in human MDM and AM at 24 h, we next wanted to determine if dexamethasone inhibited the acute glycolytic response of the human macrophage to Mtb. In order to control for changes in basal glycolysis and specifically assess if macrophage responses to Mtb were altered, MDM were treated with VC or dexamethasone immediately prior to placing them in the Seahorse XFe24 Analyzer. MDM were left to equilibrate for 20 minutes and then stimulated with irradiated Mtb or Seahorse medium, cells were continually monitored for 2 h after which a glycolytic rate assay was performed. There was no difference between VC or dexamethasone treated groups at the beginning of the assay (Figure 3A-B). The increase in induced glycolysis by MDM in response to Mtb stimulation is evident in the GlycoPER traces (Figure 3B) and was significantly enhanced compared with unstimulated MDM (Figure 3C). Dexamethasone prevented MDM from increasing glycolysis in response to Mtb (Figure 3C). Interestingly, in unstimulated MDM dexamethasone impaired glycolysis, suggesting steroids are capable of rapidly restricting metabolism (Figure 3C). Even when oxidative phosphorylation was inhibited and MDM were forced to utilize compensatory glycolysis, Mtb stimulated MDM had enhanced levels of glycolysis compared with unstimulated controls (Figure 3D). Dexamethasone inhibited compensatory glycolysis in both unstimulated and Mtb stimulated MDM (Figure 3D). Similar to treating MDM with dexamethasone for 24 h, energy maps depicting MDM treated acutely with dexamethasone displayed a leftward horizontal shift, indicating a less glycolytic phenotype, in both unstimulated (Figure 3E) or Mtb stimulated MDM (Figure 3F). Since no difference in OCR was observed in energy maps, a MitoStress test was employed to measure mitochondrial metabolism in dexamethasone treated macrophages. Mitochondrial metabolism was not altered by dexamethasone in either unstimulated or Mtb stimulated MDM (Figure S2).

**Figure 3:**
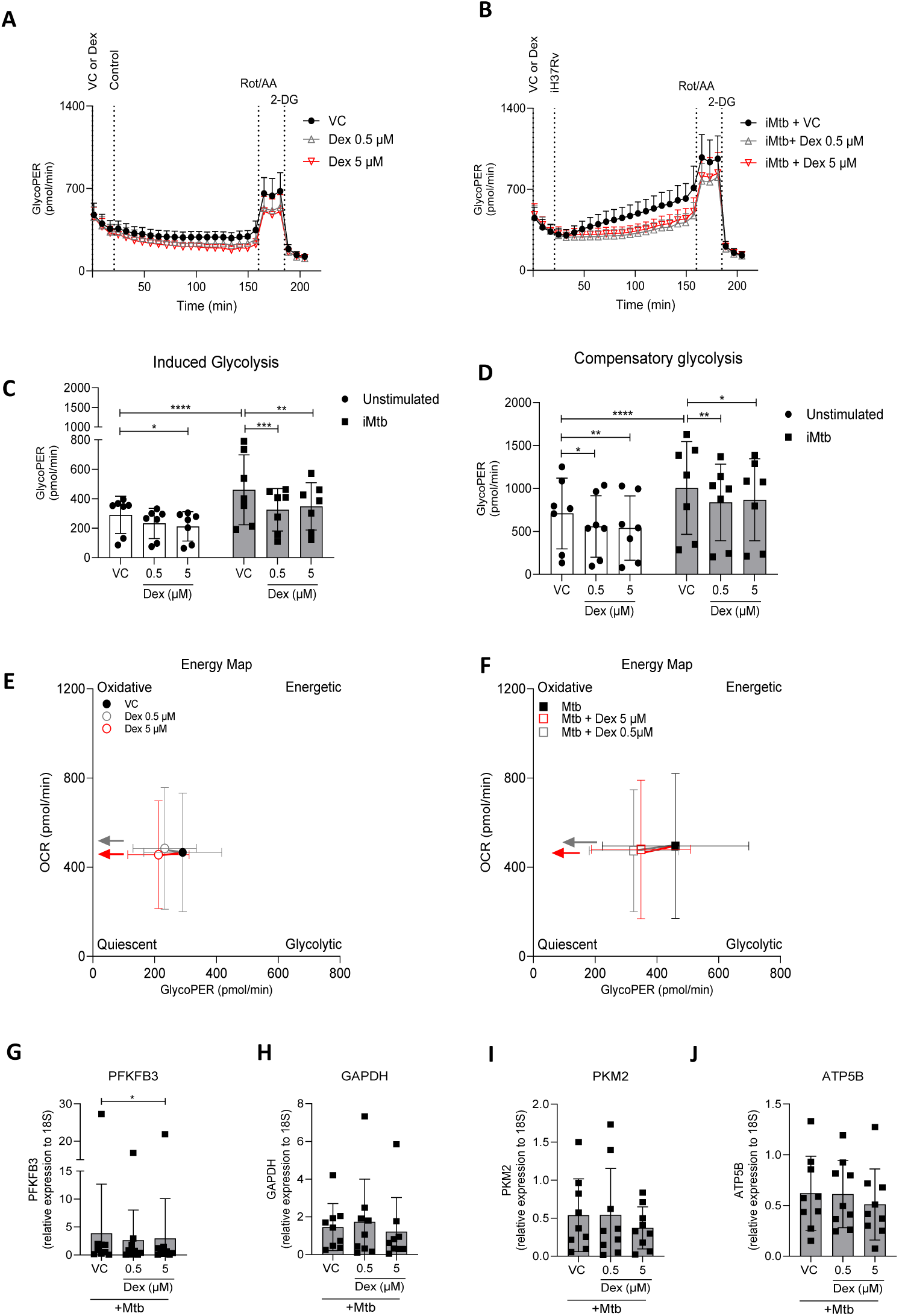
Dexamethasone impaired the glycolytic response to Mtb in human MDM. PBMC were isolated from buffy coats and MDM were adherence purified for 7 days in 10% human serum. MDM were cold lifted and plated on Seahorse XFe24 (Agilent) culture plates (A-F). MDM were treated with VC (ethanol 0.02%) or dexamethasone (0.5 or 5 µM) right before transferring cells into a Seahorse XFe24 Analyzer. After 20 minutes MDM were stimulated through the ports of the Analyzer with control (A; Seahorse media) or irradiated Mtb (B; iMtb; strain H37Rv, MOI 1-10). ECAR and OCR readings were recorded at regular intervals for 150 minutes. After 150 minutes a glycolytic rate assay was performed by injection of Rotenone/Antimycin A followed by 2DG as indicated to calculate GlycoPER from which the induced (C) and compensatory glycolysis (D) were determined. Energy maps were generated from induced glycolysis and the corresponding OCR values (E,F). Cell density was corrected for using Crystal Violet for Seahorse assays (A-F). Data are represented as means ± standard deviation. Statistically significant differences were determined using a two-way ANOVA with a Tukey post-test (C,D; n=7) or Friedman test with Dunn’s post-test (G-J; n=9); *p≤0.05, ** p≤0.01, *** p≤0.001, **** p≤0.0001.

As we previously demonstrated that dexamethasone inhibited PFKFB3 (Figure 1G) expression in MDM, we next examined whether dexamethasone reduced the expression of rate-limiting glycolytic enzymes in MDM after Mtb infection. MDM were treated with VC or dexamethasone (0.5 or 5 μM) for 1 h prior to infection with Mtb, 3 h after infection the mRNA expression of *PFKFB3, GAPDH, PKM2* and *ATP5B* was assessed by real-time PCR. Consistent with the previous observation in unstimulated MDM dexamethasone significantly reduced the expression of *PFKFB3* in MDM infected with Mtb (Figure 3G). Expression of *GAPDH, PKM2* and *ATP5B* was not altered in MDM treated with dexamethasone (Figure 3H-J).

### Dexamethasone reduces cytokine secretion by human macrophages

Changes in macrophage metabolism are associated with altered cytokine production (16–18) and it has been previously described that glycolysis is essential in regulating the secretion of both IL-1β and IL-10 in response to Mtb or LPS (10, 13, 14, 16, 19, 20). Glucocorticoids such as dexamethasone are known to reduce the production of pro-inflammatory cytokines. Since dexamethasone significantly reduced glycolysis in human macrophages, we wanted to determine if this reduction also corresponds to a decreased ability to produce cytokines.

Human MDM were treated with dexamethasone (0.05, 0.5 or 5 μM) or VC for 1 h prior to infection with Mtb (MOI 1-10). Supernatants were harvested 72 h post infection and the secretion of IL-1β, TNF, IL-6, IL-8, and IL-10 was quantified by ELISA. Mtb infection of human MDM led to an increase in the secreted levels of all cytokines assayed (Figure 4A-E). Dexamethasone significantly decreased the secretion of the inflammatory cytokines IL-1β, TNF, IL-6 and IL-8 (Figure 4A-D). The anti-inflammatory cytokine IL-10 was also dose dependently decreased by dexamethasone (Figure 4E).

**Figure 4:**
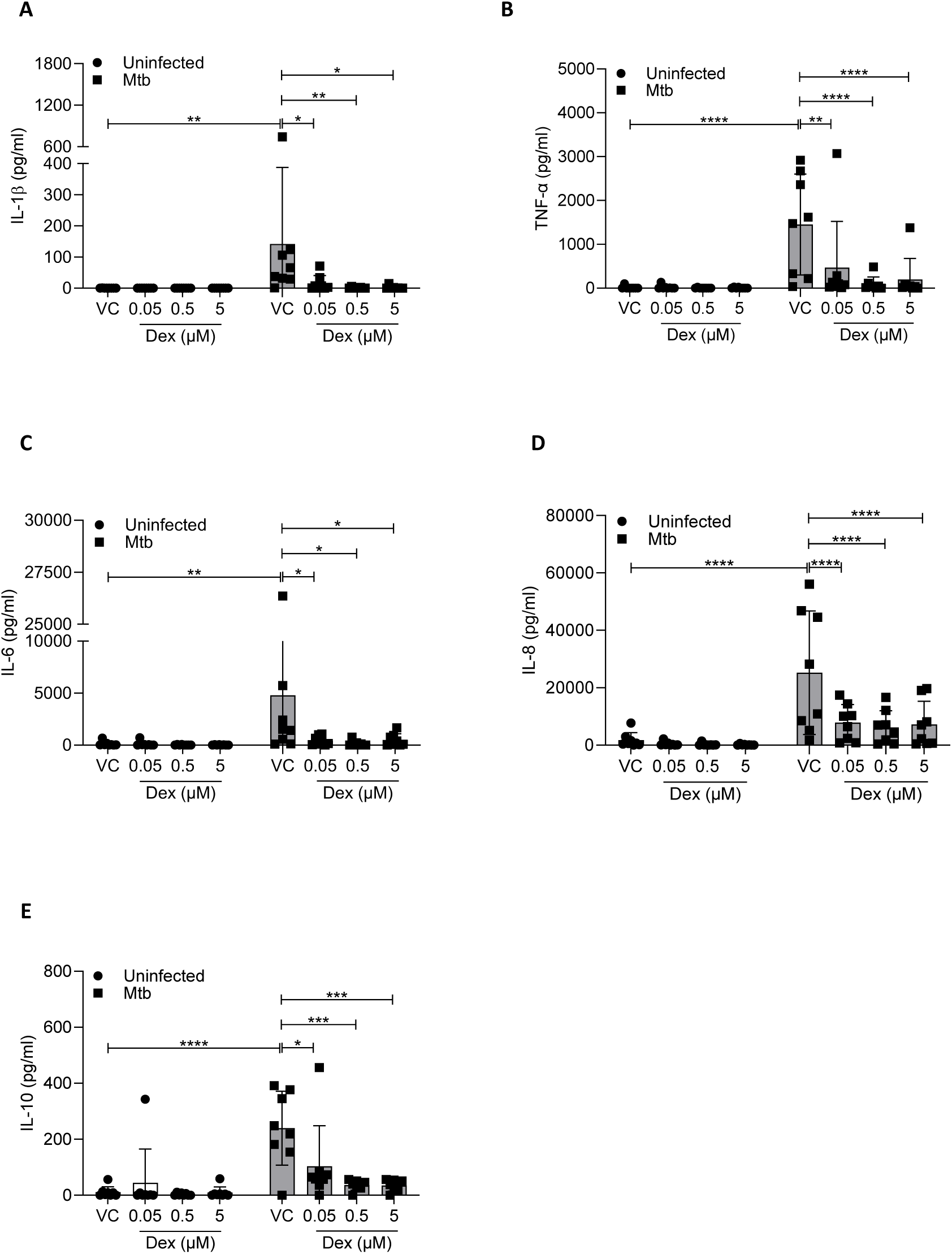
Dexamethasone reduces the production of cytokines in human MDM infected with Mtb. PBMC were isolated from buffy coats and MDM were adherence purified and differentiated for 7 days in 10% human serum. MDM were treated with VC (ethanol 0.02%) or dexamethasone (0.05, 0.5 or 5 μM) for 1 hour prior to infection with Mtb (MOI 1-10). Supernatants were harvested 72 hours post infection and the secretion of IL-1β (A), TNF-α (B), IL-6 (C), IL-8 (D) and IL-10 (E) was quantified by ELISA. Data are represented as means ± standard deviation. Statistical significance was determined using a two-way ANOVA with a Dunnett’s post-test *p≤0.05, ** p≤0.05, *** p≤0.005, **** p≤0.001 (n=8).

Dexamethasone has been reported to induce apoptotic cell death in human monocytes (21) whereas Mtb infected murine macrophages treated with dexamethasone had improved cellular survival (22). Since dexamethasone inhibited all cytokines by MDM in response to Mtb infection, it was therefore necessary to examine the effect of dexamethasone on cell viability of Mtb infected human MDM. Human MDM were treated with dexamethasone (0.05, 0.5 or 5 μM) or VC for 1 h prior to infection with Mtb (MOI 1-10) and cell death was assessed by PI inclusion assay. Mtb infection significantly increased the percentage of PI positive MDM, which was significantly reduced when MDM were treated with dexamethasone (Figure 5A). To more specifically examine apoptosis the CellEvent™ Caspase-3/7 assay was employed. Mtb infection increased the percentage of cells staining positive for activated caspase 3/7. Dexamethasone treatment completely inhibited the activation of caspase 3/7 in response to Mtb (Figure 5B). Since dexamethasone completely inhibited caspase 3/7 activation in MDM, but PI staining was more modestly reduced we therefore examined the effect of dexamethasone on a classic inducer of apoptotic cell death. MDM were treated with dexamethasone for 1 h prior to stimulation with staurosporine (1 μM) for 24 h. Staurosporine significantly increased cell death in MDM, which was further enhanced by treatment with dexamethasone (Figure 5C). Cumulatively, these data suggest that dexamethasone is specifically improving survival of MDM infected with Mtb.

**Figure 5:**
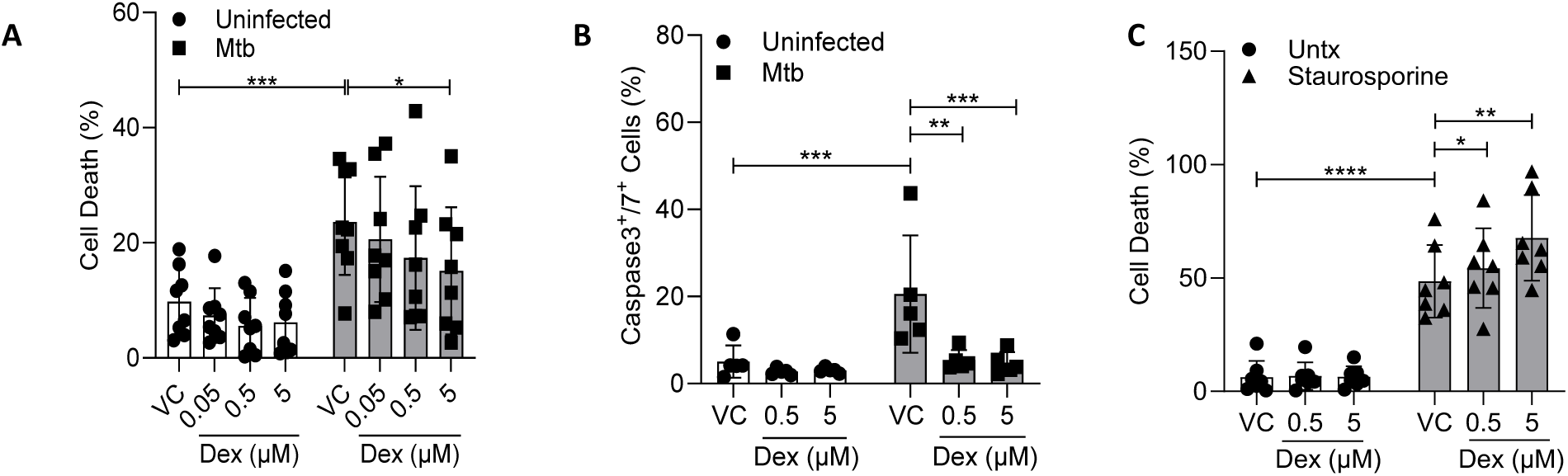
Dexamethasone reduces cell death in human MDM infected with Mtb. PBMC were isolated from buffy coats and MDM were adherence purified and differentiated for 7 days in 10% human serum. MDM were treated with VC (ethanol 0.02%) or dexamethasone (0.05, 0.5 or 5 μM as indicated) for 1 h prior to infection with Mtb (A-B; MOI 1-10, black). Alternatively, 1 h post dexamethasone MDM were left unstimulated (white) or stimulated with staurosporine (C; 1μM, grey) for 24 h, respectively. Macrophages were stained with propidium iodide (A,C) or CellEvent™ Caspase-3/7 Green Detection Reagent (B). Cells were visualized and data was analyzed using a Lionheart® FX automated microscope. Data are represented as means ± standard deviation. Statistical significance was calculated using a two-way ANOVA with Dunnett’s post-test (*p≤0.05, ** p≤0.005; A n=9, B n=5, C n=7).

### Dexamethasone restricts Mtb growth in primary human macrophages

Dexamethasone is the only approved HDT for Mtb yet the direct effect of dexamethasone on the ability of human macrophages to control Mtb growth remains unknown. Indeed, murine studies on J774 macrophages demonstrates that dexamethasone improved host cell survival, yet had no influence on intracellular Mtb growth (22). Since dexamethasone improved human MDM viability during Mtb infection in vitro, we next sought to examine the growth of Mtb in dexamethasone treated human MDM. Human MDM were treated with dexamethasone (0.5 or 5 μM) or VC for 1 h prior to infection with Mtb (MOI 1-5). MDM were then lysed at 3, 48 and 120 h post infection, plated on Middlebrook 7H10 agar supplemented with OADC and CFU were enumerated after 3 weeks (Figure 6A). There was no significant difference in phagocytosis of bacteria by Mtb 3 h post infection (Figure S3A). At 120 h post infection MDM treated with dexamethasone had a dose dependent reduction in CFU compared with control (Figure 6B). We have previously demonstrated that lactate reduces inflammatory cytokines and glycolysis, and yet improves Mtb clearance by promoting autophagy (14). Autophagy is a crucial strategy used by macrophages to combat infection, however, Mtb has been shown to target autophagic flux for survival and persistence in macrophages (23). In order to examine if dexamethasone may be limiting mycobacterial replication through autophagic induction, CFU assays were carried out in the presence and absence of bafilomycin A1, an inhibitor of the vacuolar H^+^-ATPase (V-ATPase) which inhibits phagosomal acidification. Human MDM were treated with bafilomycin A1 (50 nM) for 1 prior to treatment with dexamethasone (0.5 μM), followed by a CFU assay. There was no difference in phagocytic ability of MDM on the day of infection (Figure S3B). Bafilomycin A1 significantly enhanced mycobacterial growth in the Mtb infected human MDM (Figure 6C), even in the presence of dexamethasone. Furthermore, bafilomycin abrogated the dexamethasone mediated reduction in CFU suggesting that dexamethasone was phagolysosomal maturation to control intracellular growth of mycobacteria. We then hypothesized that dexamethasone may be inducing autophagy thereby facilitating the fusion of phagosomes with mature lysosomes to control the growth of Mtb. To test this hypothesis, MDM were treated with rapamycin (250 ng/ml), an mTOR inhibitor which promotes autophagy, for 24 h prior to treatment with dexamethasone (0.5 μM) and followed by infection with Mtb. All MDM had broadly similar phagocytic capacity on day 0 regardless of treatment (Figure S3D-E). While dexamethasone and rapamycin both reduced mycobacterial growth, treatment of MDM with rapamycin prior to dexamethasone did not produce any additive effect on reducing CFU (Figure 6D), suggesting that dexamethasone may compete with rapamycin. Cumulatively these data suggest that dexamethasone is improving the ability of human macrophages to limit Mtb growth via a mechanism that is at least in part mediated by autophagy.

**Figure 6:**
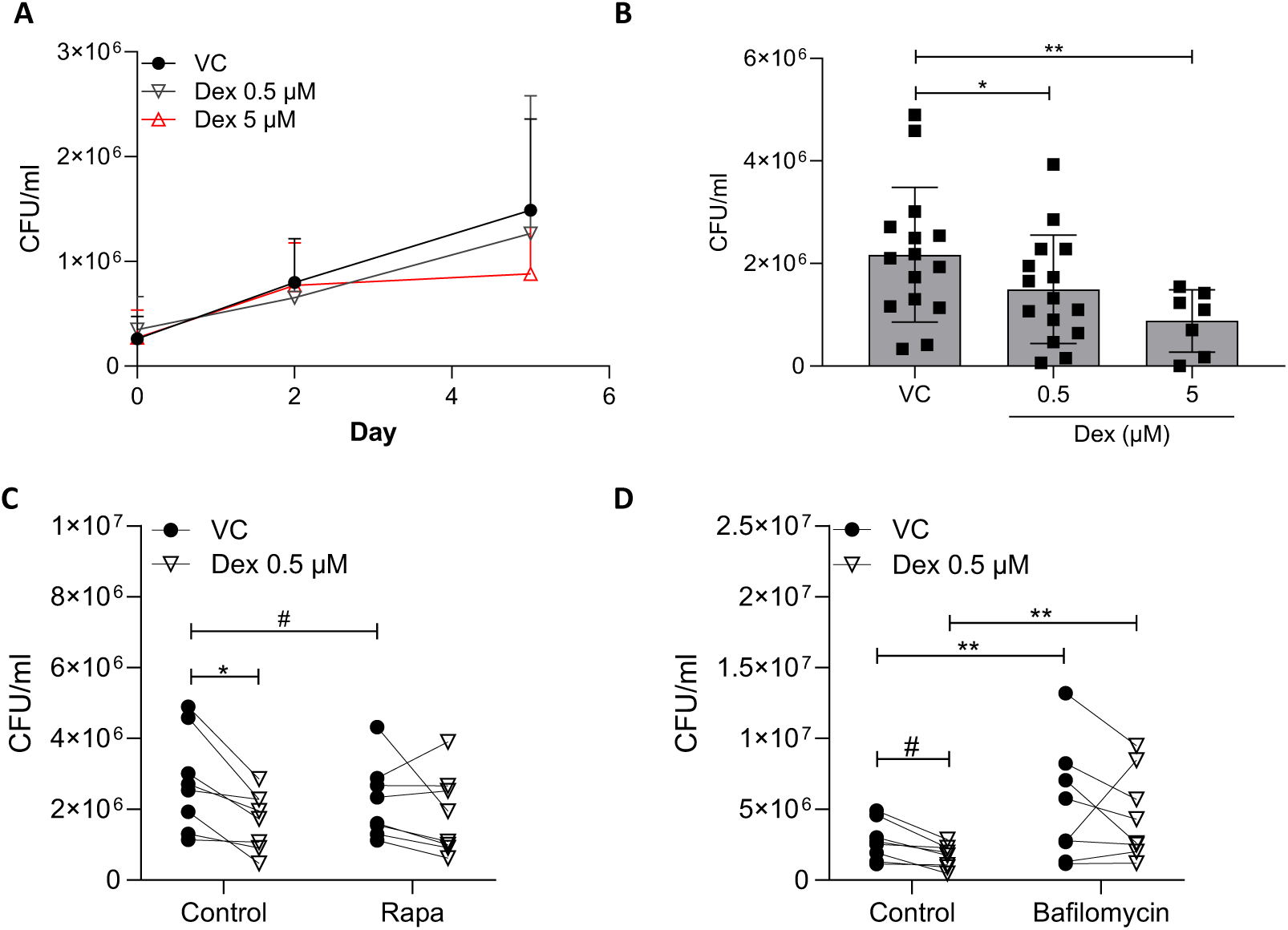
Dexamethasone controls Mtb growth in human MDM. PBMC were isolated from buffy coats and MDM were adherence purified for 7 days in 10% human serum. MDM were treated with dexamethasone for 1 hour prior to infection with Mtb (MOI 1-10). MDM were lysed and plated at 3, 48 or 120 h post infection on Middlebrook 7H10 agar supplemented with OADC (A-B). Alternatively, MDM were treated with bafilomycin A1 (50 nM) for 1 hour (C) or rapamycin (250 ng/ml) for 24 h (D) prior to dexamethasone, infected with Mtb and lysed at 120 h. Data are represented as means ± standard deviation. Statistically significant differences were determined using a one-way ANOVA with Dunnett’s post-test (B, n=8-17) or two-way ANOVA with a Holm-Sidak post-test (C-D, n=8) (*p≤0.05, ** p≤0.01) or student’s t-test (#p≤0.05).

## Discussion

GC have long been recognized for their ability to alter metabolism in organs and tissues of mammals however, the specific effects of GC on individual cell types have only started to be appreciated. We and others have reported on the crucial role that metabolism, specifically glycolysis, has in the coordinating the functional responses of human macrophages (10, 16, 20, 24). Our current study demonstrates that dexamethasone impairs glycolytic metabolism in human tissue-resident macrophages, as well as blood derived MDM. Previous observations have demonstrated that dexamethasone reduces glycolysis in stimulated murine bone marrow derived macrophages (1–3) and human MDM (3). These previous studies have utilized the extracellular acidification rate (ECAR) to describe the changes occurring in glycolysis and our data builds on this by examining the glycolytic proton efflux rate (glycoPER), which excludes mitochondrial acidification providing a more specific measurement of glycolysis. This has allowed us to identify that dexamethasone is significantly impairing glycolysis in human macrophages even in the absence of an immune insult. Consistent with previous reports (3), dexamethasone did not alter the oxidative metabolism in human MDM as measured by basal oxygen consumption rate (OCR). Human AM however, had significantly reduced basal OCR 24 h after dexamethasone treatment, therefore distinct differences are evident between human macrophage populations. Indeed, we have previously demonstrated that AM are more reliant on oxidative metabolism compared with MDM after IFN-γ treatment (10), this reliance on oxidative metabolism by the AM may be why it is more sensitive to treatment with dexamethasone.

The reductions in glycolysis by dexamethasone was evident in macrophages after 2 h and impaired the ability of the MDM to induce glycolysis in response to Mtb. This was also associated with the downregulation of *PFKFB3* a gene encoding a rate-limiting enzyme at the beginning of the glycolytic pathway, thereby restricting glucose utilization. The metabolic rewiring of LPS stimulated murine macrophages by dexamethasone has recently been proposed to be mediated by production of the intermediate metabolite, itaconate (1). While we hypothesized this may be occurring in our cellular model, we did not detect itaconate in MDM treated with dexamethasone, either uninfected or infected with Mtb (data not shown). Furthermore, while LPS stimulated murine macrophages had improvements in mitochondrial metabolism when treated with dexamethasone (1), mitochondrial metabolism in our human MDM model was stable in the presence of dexamethasone. While we cannot rule out that the differences in macrophage response may be due to the activating stimulant, it strengthens previous reports that mitochondrial dynamics differ fundamentally between human and murine macrophages (10, 13, 15, 25).

Glycolytic metabolism in macrophages has been intrinsically linked to cytokine production (26–28), specifically IL-1β, production and secretion by macrophages (10, 12, 16, 17, 19, 29) and human monocytes treated with dexamethasone have reduced IL-1β production in response to LPS (30). While GC are broadly accepted to inhibit production of inflammatory mediators, we demonstrate for the first time that cytokine secretion was significantly inhibited in dexamethasone treated human MDM infected with Mtb, similar to dexamethasone treated human AM infected with other pathogens (31). Thus far, primary microglia are the only other human macrophages where dexamethasone has been shown to inhibit cytokine production during Mtb infection (32). While it is unsurprising that pro-inflammatory cytokines were significantly inhibited by dexamethasone, IL-10 secretion was also significantly repressed. This may provide support that the general suppression of cytokines is associated with the reduction in macrophage metabolism. We and others have previously shown that blocking glycolysis with 2-deoxyglucose (2DG) impairs secretion of both IL-1β and IL-10 in human macrophages (10, 15). Furthermore, inhibition of oxidative phosphorylation by oligomycin restored the ability of dexamethasone treated BMDM to transcribe TNF, which was not evident if HIF-1α was pharmacologically or chemically induced (2). These data cumulatively suggest that metabolism may be an essential mechanism targeted by GC in macrophages to limit cytokine production.

GC are known to act as both inducers and inhibitors of cell death pathways such as apoptosis and necrosis, which differs depending on cell type, species and environmental milieu (22, 33–35). Dexamethasone increased cell death of MDM induced by staurosporine, however, dexamethasone reduced cell death in human MDM infected with Mtb, which was associated with less caspase activation. In line with our observations, previous work has identified that dexamethasone inhibits cell death pathways in both human and murine Mtb-infected macrophages (22), linked to preventing the dissociation of the hexokinase II from the mitochondrial membrane. Human lung fibroblasts were also protected from Mtb induced cell death by dexamethasone and other GC (34). Moreover, GC and not mineralocorticoids improved the ability of human MDM to control intracellular Mtb growth, which was not replicated in the absence of macrophages (34). This suggests that GC improve the ability of macrophages to limit Mtb replication. While other studies in murine macrophage cell lines have shown no difference in Mtb growth after dexamethasone treatment (22) or a reduced ability to control other mycobacterial infections (36), here we provide further evidence that dexamethasone improves the ability of the human MDM to restrict intracellular Mtb. We propose that this improved ability to limit replication is at least in part mediated by enhanced macrophage survival and autophagy. Indeed, we have previously shown that IL-10 production is mediated by glycolysis (10), and that IL-10 can inhibit phagosome maturation allowing Mtb to replicate (37). Our current work demonstrates that both glycolysis and IL-10 are reduced by dexamethasone and that this is concurrent with improved bacillary control, suggesting that early IL-10 production may critically influence infection outcomes in human macrophages.

## Limitations of the current study

We acknowledge that the current study is a reductive cellular model of Mtb infection and proposes beneficial effects on macrophages in the absence of other immune cells. Reductive cellular models of infection and steroid treatment are necessary to decipher the intricate interactions in individual immune cell populations to optimize their clinical utility for patient treatment. Our cellular data supports a cell-targeted therapeutic approach and will facilitate the development of *in vivo* preclinical models and clinical trials for future macrophage-focused therapies.

While we demonstrate that dexamethasone improves survival of Mtb infected macrophages and separately show that dexamethasone treated macrophages have reduced CFU. We do not have mechanistic evidence to causally associate enhanced cellular survival and enhanced bacterial clearance, however, we postulate that these two observations are associated. Further studies are required to determine whether dexamethasone enhancing macrophage survival improves bacterial clearance, or whether improved bacterial clearance in dexamethasone treated macrophages contributes to increased survival.

GC, such as dexamethasone, are a mainstay of treatment when patients with TB present with paradoxical reactions (PR) (38). While the etiology of PR in patients who have been previously treated for TB is unknown, it is likely caused by an excessive immune reaction to dead and/or lingering unculturable bacilli, or to persisting antigens (39, 40). The current study provides further rationale that treating these patients with GC could limit inflammation by decreasing cellular metabolism resulting in reduced cytokine production and helping to promote clearance of the bacilli and persisting antigens.

These data provide *in vitro* evidence to further investigate the use of GC as an adjunctive cell-targeted host directed therapy for TB and other respiratory infectious diseases by influencing the metabolism and function of human macrophages. These results support the development of targeting GC therapies to macrophages thereby mitigating inflammation without compromising ancillary host antimicrobial defense and offer a rationale for the clinical benefit of GC in some TB presentations.

## Acknowledgements

We would like to acknowledge the Irish Blood Transfusion Service for supporting our research by approving us to use anonymized un-transfused blood components for our research. We gratefully acknowledge all people undergoing bronchoscopy at St. James’s Hospital Dublin who consented to take part in our research. We acknowledge the key contributions of the Clinical Research Facility at St. James’s Hospital and the bronchoscopy suite, and the core facilities at the Trinity Translational Medicine Institute. The following reagent was obtained through BEI Resources, NIAID, NIH: Mycobacterium tuberculosis, Strain H37Rv, Gamma-Irradiated Whole Cells, NR-49098.

