## Supplementary figures and images for "Dexamethasone impairs glycolysis but improves mycobacterial killing in primary human macrophages"

### Figure S1

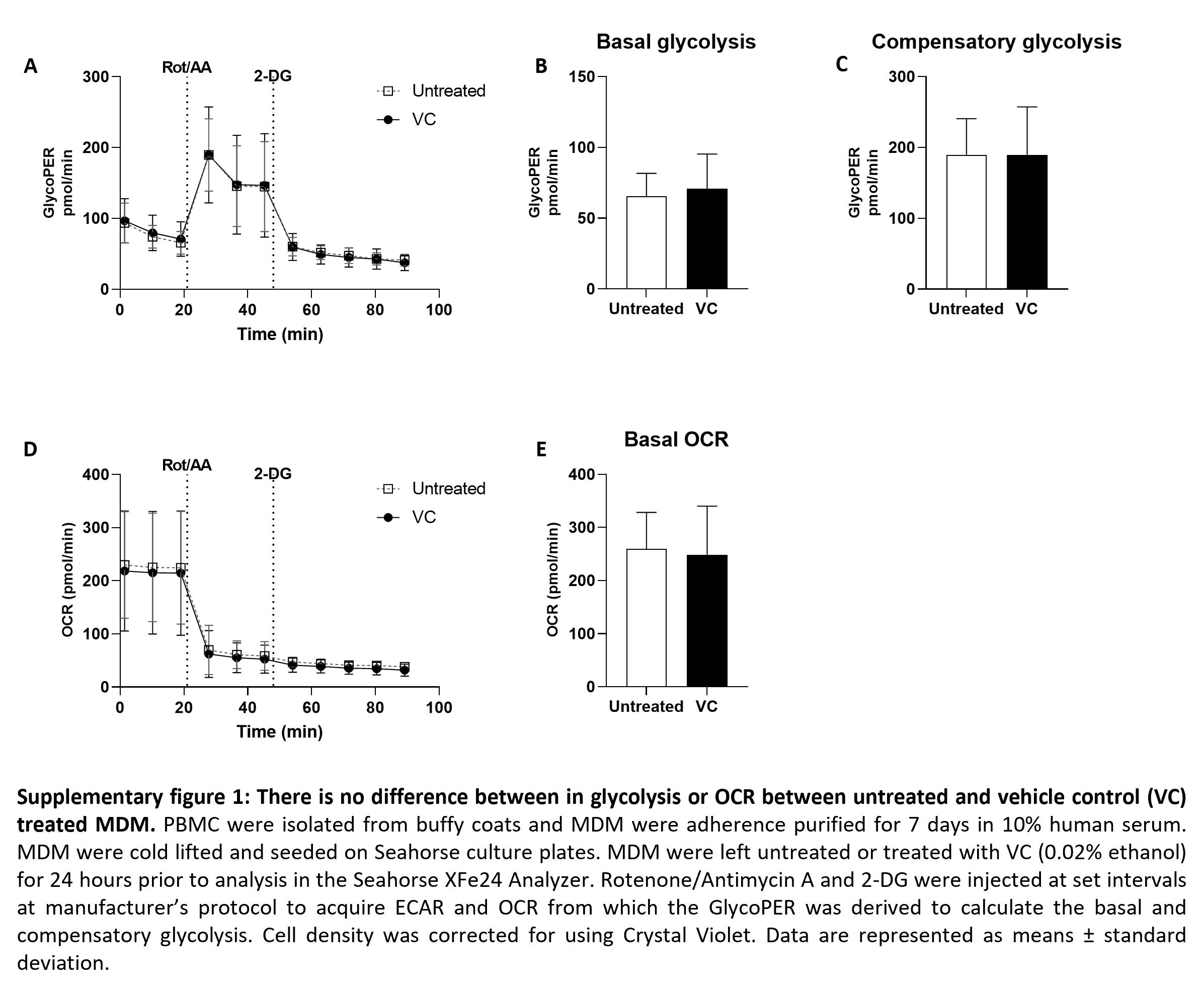

### Figure S2

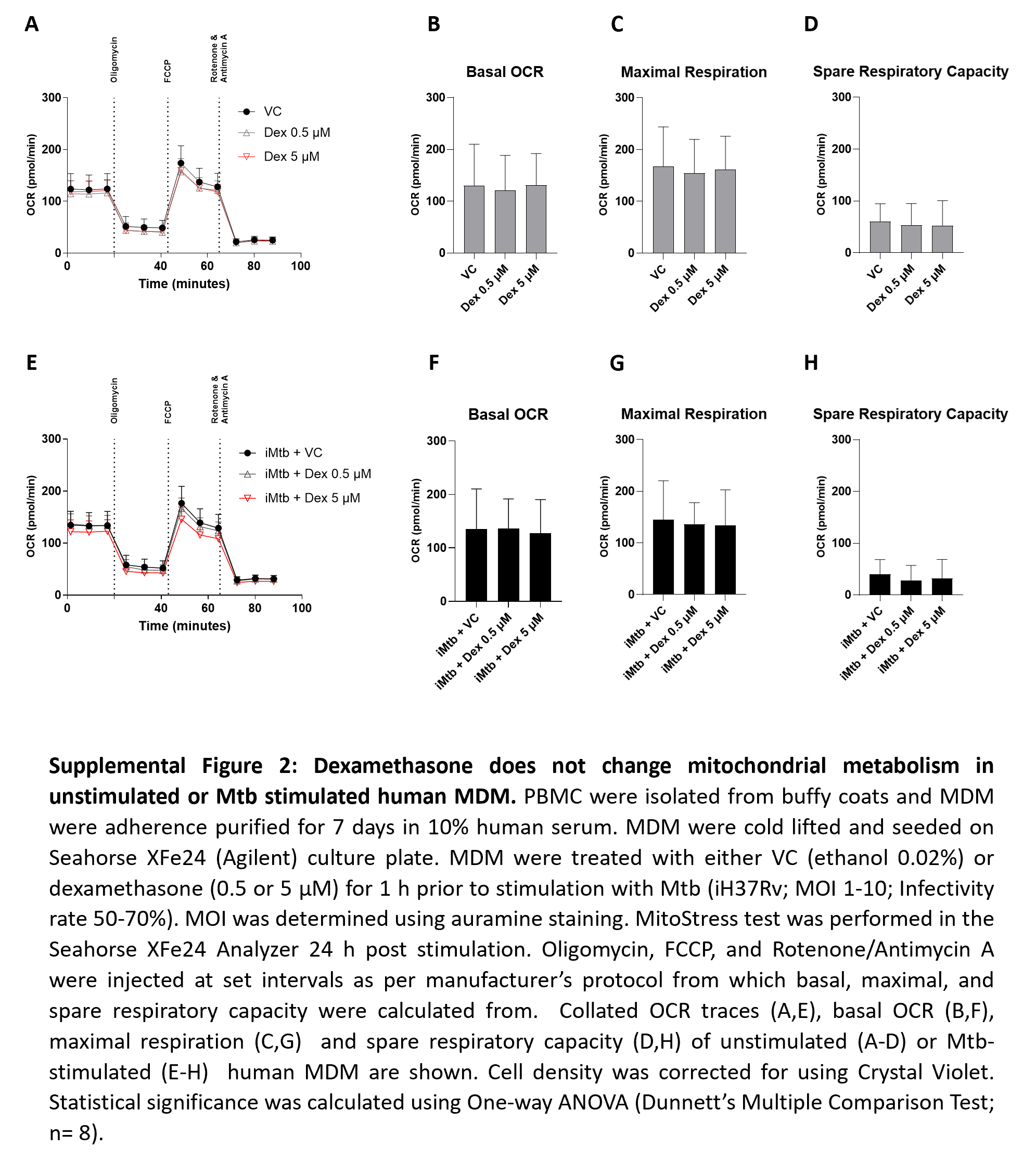

### Figure S3

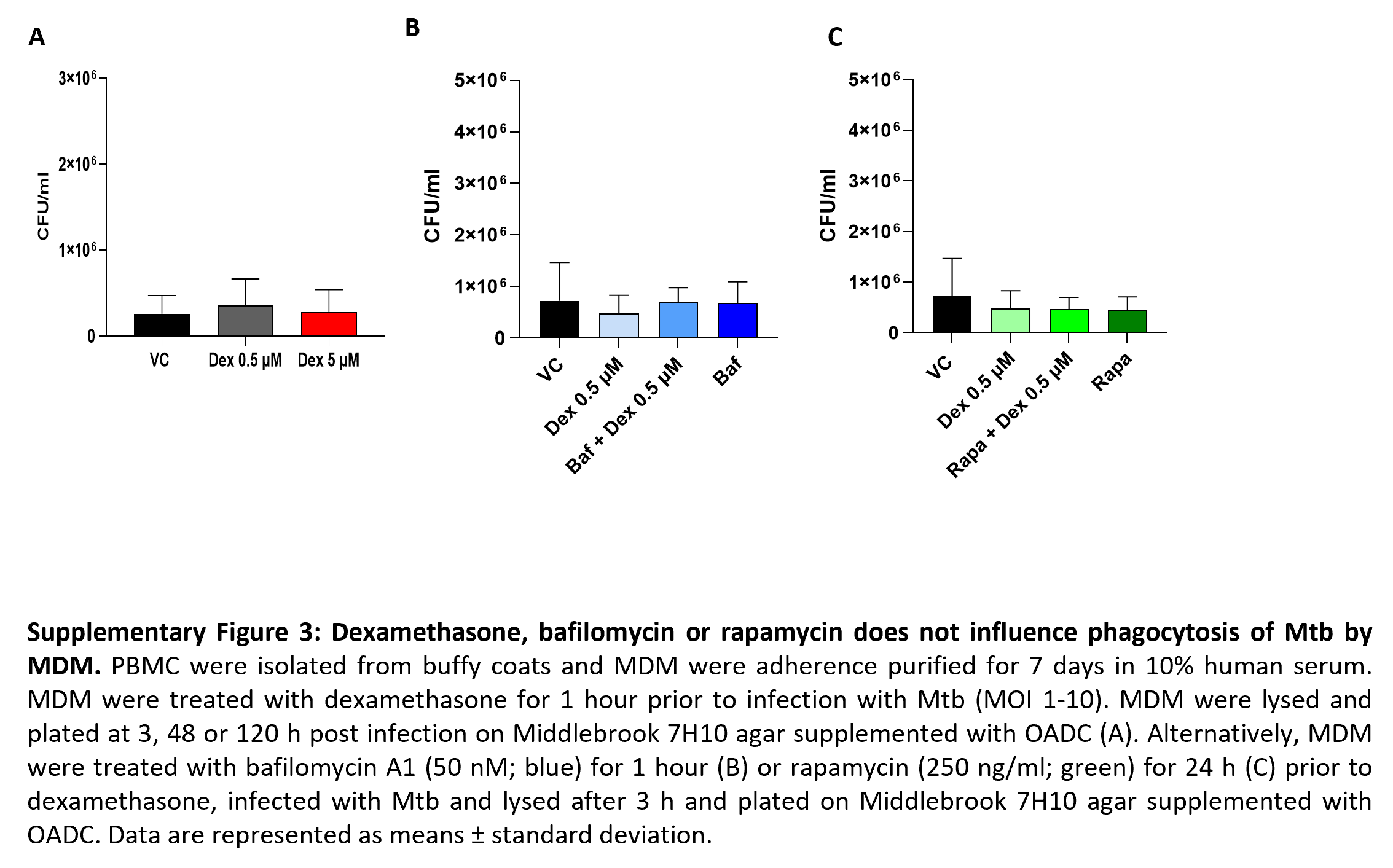
